# Automated non-lethal moth traps can be used for robust estimates of moth abundance

**DOI:** 10.1101/2022.06.13.495870

**Authors:** Jonas Mielke Möglich, Patrick Lampe, Mario Fickus, Jannis Gottwald, Thomas Nauss, Roland Brandl, Martin Brändle, Nicolas Friess, Bernd Freisleben, Lea Heidrich

## Abstract

1. Recent reports of insect decline highlight the need for extensive large-scale insect monitoring. However, obtaining abundance or species richness data at high spatial and temporal resolution is difficult due to personnel, maintenance, and post-processing costs as well as ethical considerations. Non-invasive automated insect monitoring systems could provide a solution to address these constraints. However, every new insect monitoring design needs to be evaluated with respect to reliability and bias based on comparisons with conventional methods.
2. In this study, we evaluate the effectiveness of an automated moth trap (AMT), built from off-the-shelf-hardware, in capturing variations in moth abundance, by comparing it to a conventional, lethal trap. Both trap types were operated five times on 16 plots from the beginning of July 2021 to the end of August 2021.
3. Moth abundance scaled isometrically between the two trap types. Consequently, the respective seasonal patterns in abundance determined over the monitoring period were similar.
4. The AMT samples phenological patterns using a robust and non-lethal method. However, an initial quantitative in-field test revealed that its long-term applicability must be preceded by several adjustments to the power supply and to data transfer. Depending on the software implementation, the AMT can be used to address a broad range of research questions while also reducing both energy expenditure and the disturbance of non-target animals.

## Introduction

Recent reports on the decline in insect abundance and biomass (e.g., Hallmann et al., 2017, 2020; Seibold et al., 2019) have raised public concern as well as calls for immediate conservation actions (Harvey et al., 2020). However, shortcomings in data availability have cast doubt on the generalizability of this decline (Eggleton, 2020; Simmons et al., 2019), while highlighting the need for regular and comprehensive assessments of insect communities (Didham et al., 2020; Hausmann et al., 2020; Montgomery et al., 2020; Wagner, 2020). Such efforts have been hindered by the difficulty of establishing intensive monitoring programs with broad spatial and taxonomic coverage (Montgomery et al., 2021). Visual surveys such as transect walks, net sweeping or hand catches are labor intensive. Conventional traps can reduce the demands of field work, but they are limited by their high maintenance and post-sampling efforts, their considerable expenses, and their logistical challenges (Montgomery et al., 2021). Furthermore, caught specimens must be stored at least until their identification, but storage or freezer space may not be readily available. Lastly, conventional insect monitoring methods usually involve killing the insects for subsequent determination and biomass assessment, which has raised ethical concerns, because the design of some traps can result in the slow death of the insects that must be weighed against the knowledge and conservation benefits gained from the acquired data. The debate over consciousness of insects and their ability to feel pain should be recognized (Fischer & Larson, 2019). Even in the absence of detrimental effects imposed by the long-term monitoring of insect populations (Gezon et al., 2015), less destructive alternatives should be considered whenever possible (Drinkwater et al., 2019). Consequently, new, non-lethal monitoring systems are required that can be readily deployed in insect surveys.

Given the above considerations, most monitoring campaigns focus on specific taxa or locations, with the latter often being chosen based on accessibility and the high density of the target taxon (Didham et al., 2020; Fournier et al., 2019). However, this could result in site selection bias and thus to false trend estimations (*missing zero effect*, (Didham et al., 2020)).

Another practice is to focus on sampling periods that are assumed to adequately cover insect diversity (Ozaki et al., 2011; Scalercio et al., 2012). However, such temporal snapshots, whether based on daily or on annual data collection, imply a loss of information (Ozaki et al., 2011), because species occurring outside the phenology peak will remain undetected, as will phenological shifts (*snapshot* & *groundhog effect*, (Didham et al., 2020)). Monitoring networks that collect data on a daily basis, such as the Rothamsted insect survey (*The Insect Survey*, 2021), are a notable but rare exception. Yet even this dataset, made up of > 10 million records, and thus probably more accurate than any other, cannot provide complete species inventories at the spatial resolution relevant for conservation (Sánchez-Fernández et al., 2021).

Automation can reduce both the work load and the cost associated with monitoring, in turn facilitating the collection and identification of insects at high temporal and spatial resolution (Didham et al., 2020; Montgomery et al., 2021). The rapid development of remote sensing and data acquisition (e.g., Müller & Brandl, 2009), especially sensor-based systems, but also those based on radar, acoustic monitoring, or image acquisition (Høye et al., 2021; Noskov et al., 2021), offer new options in biological studies.

Prior to their wider adoption, newly designed trap types need to be evaluated and compared with conventional methods (assuming they provide reliable results), because every design change affects capture efficiency (Preti et al., 2021) and therefore conclusions regarding species composition. For example, flight interception traps consisting of only one collecting jar at the bottom, rather than an additional collecting jar at the top, might underestimate taxa that tend to move upwards after collision (Knuff et al., 2019). Even in traps targeting single species, slight adjustments such as the width of the trap opening or the addition of a rain cover can affect the number of captured individuals (Burner et al., 2021; Guarnieri et al., 2011). In extreme cases, design-related differences in the community and/or abundance data can bias ecological conclusions (Saunders & Luck, 2013) and thus impact conservation or management measures. It is therefore necessary not only to compare the efficiency of newly designed traps with that of conventional traps, but also to ask whether the former are able to accurately reflect the response of the target group to changes in the environment.

Moths are one of the most diverse insect groups and they are closely tied to their ecosystems (Fox, 2013). Accordingly, they have a long history in monitoring studies (Sánchez-Fernández et al., 2021) and nowadays are often used as model organism in tests of new identification algorithms (Chang et al., 2017; Poremski, 2017; Wu et al., 2019). However, moths are highly sensitive to trap design (Brehm & Axmacher, 2006; Fayle et al., 2007; Hausmann et al., 2020). For example, differences were observed when bucket traps rather than recordings from a light sheet were used as the detection method. A light sheet is often more effective, as species are recorded directly when they land, whereas not all moths necessarily access the funnel of a bucket trap (Brehm & Axmacher, 2006; Wölfling et al., 2016).

An assessment of the number of individuals is the first step in analyzing the variation in the biodiversity patterns of insects, e.g., when a certain abundance is necessary to support viable populations (Gaston, 2000). In this study, we investigate whether an automated moth trap (AMT), which attract moths to a screen via ultraviolet (UV) light and then photographs them (e.g., Bjerge et al., 2021; Hogeweg et al., 2019), can depict the same abundance trends as using conventional, lethal light traps. To the best of our knowledge, our AMT is the first automated light trap tested for moth monitoring and directly compared to a conventional lethal light trap. In addition, the AMT was tested under realistic in-field conditions, in a relatively remote area, to assess its handleability but also its weaknesses. Specifically, the AMT and a conventional bucket trap were used to assess moth abundances at the same sites throughout late summer, shortly after the phenological peak of these insects. If the two trap types are equally efficient in capturing local moth abundance, the temporal decline in individuals should not differ between them.

## Material & Methods

### Study sites

Our study is part of the Nature 4.0 framework (www.natur40.org) and was conducted in the “Marburg Open Forest”, a 1.5 km^2^ university-owned forest area near Caldern, in Hesse, Germany. The forest is a typical beech-dominated managed forest embedded in an agricultural landscape. The 18 trap sites were selected from the existing Nature 4.0 sites and were evenly distributed over the research area, thus covering small-scale climatic, structural, and ecological differences. Each site includes a beech tree (50 to 120 years-old) in its center and is separated from the other sites by least 50 m.

### Conventional traps

The conventional light traps (Appendix S1) consist of a super-actinic UV light tube (12 V, 15 W, Bioform), attached over a funnel leading to a 10 l bucket. The bucket contains a closed chloroform jar from which a piece of cloth is drawn through a small hole in the lid to act as a wick, thus filling the bucket with chloroform. Pieces of egg carton are added to allow the caught moths to rest, thus also reducing stress and wing damage. Attracted moths circle around the UV lamp, spiral downwards, land in the bucket, and are killed by the chloroform. The UV light is powered by a 12 V lead battery. A light switch (Kemo Germany M197) attached to the battery automatically turns the light on at sunset and shuts the light off at sunrise. In the morning, the traps are emptied and all moths are collected and stored in a freezer for later sorting.

### Automated moth traps

A prerequisite of our AMT is that it should operate in a real-world environment and in a variety of settings. Therefore, the design requirements of the AMT are as follows:

- modular and flexible, allowing both extensibility and adaptability for further applications
- reproducible, such that the AMT could be easily assembled by other researchers
- robust, to allow continuous operation of the trap
- easy to use and configurable, to support long deployment periods while minimizing the work load

The traps are accordingly constructed following the design of Bjerge and colleagues (2021), with a few hardware modifications and using our own software. The AMT consists of a UV tube that attracts the moths towards a white, cloth-lined LED screen, on which they rest. The background illumination allows for standardized photos, which are taken by an oppositely placed camera box with an attached ring light for better illumination. The camera box consists of a Raspberry Pi computer with an attached camera. The whole system is mounted on a tripod and powered by an external battery box with a 12 V battery as well as a power bank inside the camera box. The Raspberry Pi’s software is programmed to trigger a customizable schedule, according to which photos are taken and the light (UV and LED screen) is powered. All of the components, their prices, and the measured power consumption are listed in Appendixes S2 and S3 (for AMT’s architecture and a detailed description of the trap design, see Appendix S1). To ensure design reproducibility, the trap is based on off-the-shelf hardware components.

To enable the longest possible operation time, only the UV light and LED screen are powered by the 12 V battery, so that the AMT can be left unattended in the field for up to two nights. An additional advantage of this design is that the voltage remains stable even when the LEDs are powered, since the Raspberry Pi, which is susceptible to voltage fluctuations, has its own 5 V power supply. This setup also avoids energy loss during voltage conversion. The system is modular, with a sensor box at its core that switches the UV light and the LED screen on and off by means of relays. Both the UV light and the LED screen are connected to the sensor box via waterproof cables and connectors, so that all circuitry is contained in the sensor box. The specific UV lamp or LED screen can thus be replaced without having to change the sensor box itself. Only the ring light, which is also controlled by a relay, is permanently connected to the sensor box. To ensure robustness, all components, except the ring light, are waterproof.

Ease of use is ensured by using standardized plugs and reverse-polarity-protected sockets for the individual components. In addition, the software is designed to be configurable and to start automatically.

Our software implementation is based on an extended version of the software of Gottwald et al. (2021), which was built using the *pimod* tool (Höchst et al., 2020) to configure operating system images in a user-friendly manner. Triggering the camera and the control pins of the Raspberry Pi by the program switches the external components (i.e., the ring light, LED screen, UV light) on and off via attached relays. AMT’s software is configurable by an *yml* file. A downloadable image that can be customized with *pimod* to fit the needs of the project is available in our GitHub repository (https://github.com/Nature40/InsectPhotoTrapOS/releases/tag/IPTv1.0.7). An additional feature of the software is that the sensor box includes a Wi-Fi connection, which allows wireless observation and configuration. The box can be accessed via *ssh* (Secure Shell) or via the live view of the camera using *http* (Hypertext Transfer Protocol) and a browser on the mobile device.

### Sampling/experimental design

The two trap types were tested at separate times, to avoid their mutual interference during sampling. Thus, the eight AMTs and eight conventional trap systems were randomly assigned to the 16 sites for one night. The following night, the trap types were exchanged such that sites assigned an AMT the first night received a conventional trap the following night and vice versa (two nights = one round). This routine was performed five times from July 2021 to August 2021. The AMTs were scheduled to take photos every 30 s. Activation of the lights (UV and LED screen) was scheduled to start at 10 p.m. and end at 6:00 a.m., with an on time of 50 consecutive min/h followed by 10 min of lights out. This was done to test the functionality of the hourly schedule and to roughly mimic the light phase of the conventional trap. Examples of the configuration of our specific unit and its operation schedule are shown in Appendix S4. With this schedule, the theoretical power consumption was 19.175 Wh (see Appendix S2).

### Moth counting

Moth abundance in the photos taken with the AMT was determined by counting the number of moths in each frame. Thus, in each photo, the total number of moths present on the LED screen was counted manually. For the total number of moths per night, only new arrivals on the LED screen were counted; in other words, when the total number of resting insects in a photo superseded the total number of resting insects in the prior photo, the moth count was raised. This method was used because movements of the insects on the LED screen made it difficult to identify individuals. In addition, the possibility that the same individual flew out of the photo, swirled around the light, and then again rested on the LED screen could not be ruled out. Accordingly, in the frame by frame counts of the insects, only the total number of currently sitting insects was considered. Non-target insects were not counted. Due to varying image quality, very small insects could not be identified with confidence.

Samples from conventional bucket traps were determined on the order level using a binocular, and non-target insects were excluded.

### Statistical analyses

The analyses are aimed at testing whether the number of moths captured by the AMTs scale isometrically to the number of moths captured by the conventional traps, thus yielding similar seasonal patterns in the number of individuals.

Our tests for an isometric relationship between the two traps are based on the assumption that the two nights of each round were comparable, i.e., that the number of moths do not differ when one or the other trap system was installed. Unfortunately, due to several failures of the AMTs (described and discussed below), only some of the sampled locations had repeated measures (Appendix S5). Thus, we tested *a priori* for date-wise differences between the number of moths caught by the conventional traps, which had a larger sample size due to the AMT malfunctions. This pairwise comparison, using an ANOVA followed by a post-hoc test, revealed that rounds one, three, and five had comparable numbers of individuals; in rounds two and four, the number of moths during the respective test nights differed (see Appendix S5). We therefore applied the following analyses to both the full data set and to a reduced data set in which rounds two and four were omitted.

A simple linear (ordinary least squares, OLS) model was used in a first assessment. The OLS tests whether a variable Y, in this case the number of moths detected by AMTs, depends on variable X, here, the number of moths caught by conventional traps. This assumes that conventional traps reflect a true baseline of available moths. However, it is more likely that they, and the AMTs, reflect only a fraction of the true abundance present at the time and location of trap deployment and that both traps are dependent on this temporal and spatial baseline. Thus, in a second step the major axis (MA) method was used in the analysis, as it accounts for variations in the X variable and X and Y are interchangeable (see Smith (2009) for details). The MA was calculated with 999 permutations, using the lmodel2-function of the lmodel2 package (Legendre, 2018).

While some of the plots were repeatedly sampled, which violated the assumption of independence, most of the plots were sampled only once or twice due to trap malfunctions (see Appendix S5). This hindered the implementation of PlotID as a random effect, because the resulting singularities could have led to overfitting as well as numerical problems and would have made standard inferential procedures inappropriate (Bates et al., 2015). Also, an ANOVA did not show any significant differences between the plots, suggesting that the plot had a neglectable influence. Nevertheless, we repeated the OLS and MA for the average number of moths per trap type and round. This procedure also addressed the effect of the sampling round on the number of moths, while allowing the same model parameters to be used across the full and reduced data sets as well as across the OLS and MA, with the latter not accommodating more than one explanatory variable. Note, however, that because the results of the reduced data set were based on a very low sampling size, they should be interpreted with particular care. To handle the discrete nature of the moth counts, the counts from AMTs and conventional traps were log_*e*_-transformed in all analyses. This allowed retention of the data of both axes within a comparable range (see Appendix S6 for GLMs with an untransformed dependent variable, a log_*e*_-transformed independent variable, and a negative-binomial distribution). Whether the slopes resulting from all models equaled one was determined using the *slope.test* function of the smatr-package (Warton et al., 2012).

We then tested whether the two trap systems differed in depicting abundance trends, by taking advantage of the phenology of moths. In Europe, moths usually peak in abundance and diversity around July (Roth et al., 2021; Sánchez-Fernández et al., 2021). The later the sampling time thereafter, the fewer moth individuals in the area. We thus used the Julian day, which well-reflected the time lapse between the rounds and thereby possible seasonal trends, as an independent variable to model the number of individuals using an OLS (see Appendix S6 for GLM and OLS modeling with “Round” as an ordered factor). The trap type was included as an interaction term to test for differences in the systems in depicting the temporal trend in abundance:

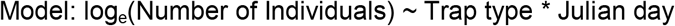

For consistency, the number of individuals was log_e_-transformed. Linear hypothesis tests were conducted using the package “car” (Fox et al., 2021). All figures were created using the package “ggplot2” (Wickham et al., 2021). All analyses were conducted using R (R Core Team, 2021) and RStudio (RStudio Team, 2021).

## Results

The conventional traps captured 17,164 individuals (mean = 226, SE = 25, min = 19, max = 885) in total and 6,755 individuals (mean = 218, SE = 36, min = 20, max = 791) considering only samples with accompanying AMTs. The latter captured a total of 4,065 individuals (mean = 131, SE = 24, min = 13, max = 616). On average, AMTs captured 87 fewer individuals (Welch two sample t-test (conducted on log-transformed data): t = −2.129, df = 59.871, p = 0.037). The variance between the two trap systems did not differ (F-test on log-transformed data F_1,30_ = 0.91, p = 0.80). In the OLS model, the relationship between the trap types in terms of moth abundance was isometric only for rounds one, three, and five (slope = 0.81, linear hypothesis test: F_1,19_ = 1.35, p = 0.26), not when all five sampling rounds were used (slope = 0.66, linear hypothesis test: F_1,29_ = 7.11, p = 0.01). The slopes of the model II regression models were close to one in all cases, and the intercepts did not significantly differ from zero (Table 1). Neither slopes modeled by the OLS nor those obtained with the model II regressions differed significantly from one when the average value per round served as the dependent variable (Table 1, Figure 1).

**Figure 1:**
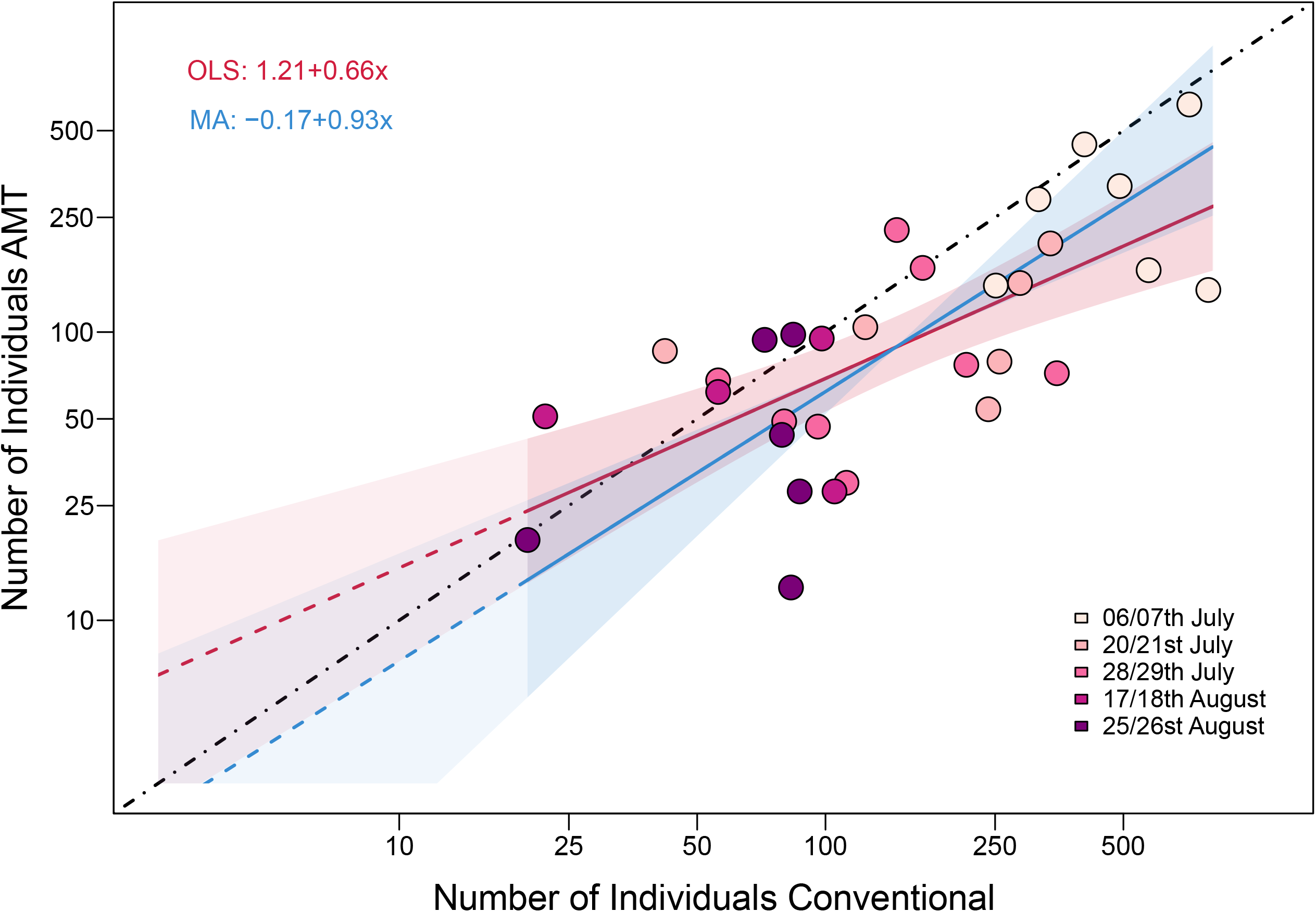
The number of individuals captured by automated moth traps (AMTs) and conventional light traps per plot and round. The points are color-coded according to the sampling dates. The black line represents an isometric relationship with an intercept of zero and a slope of one, the red line the ordinary least squares (OLS) model, and the blue line the major axis (MA) model, with shaded areas as confidence intervals. Note that the extrapolation on the left (less intensively shaded) only displays the uncertainty around the intercept. Both axes were log-transformed.

**Table 1:**
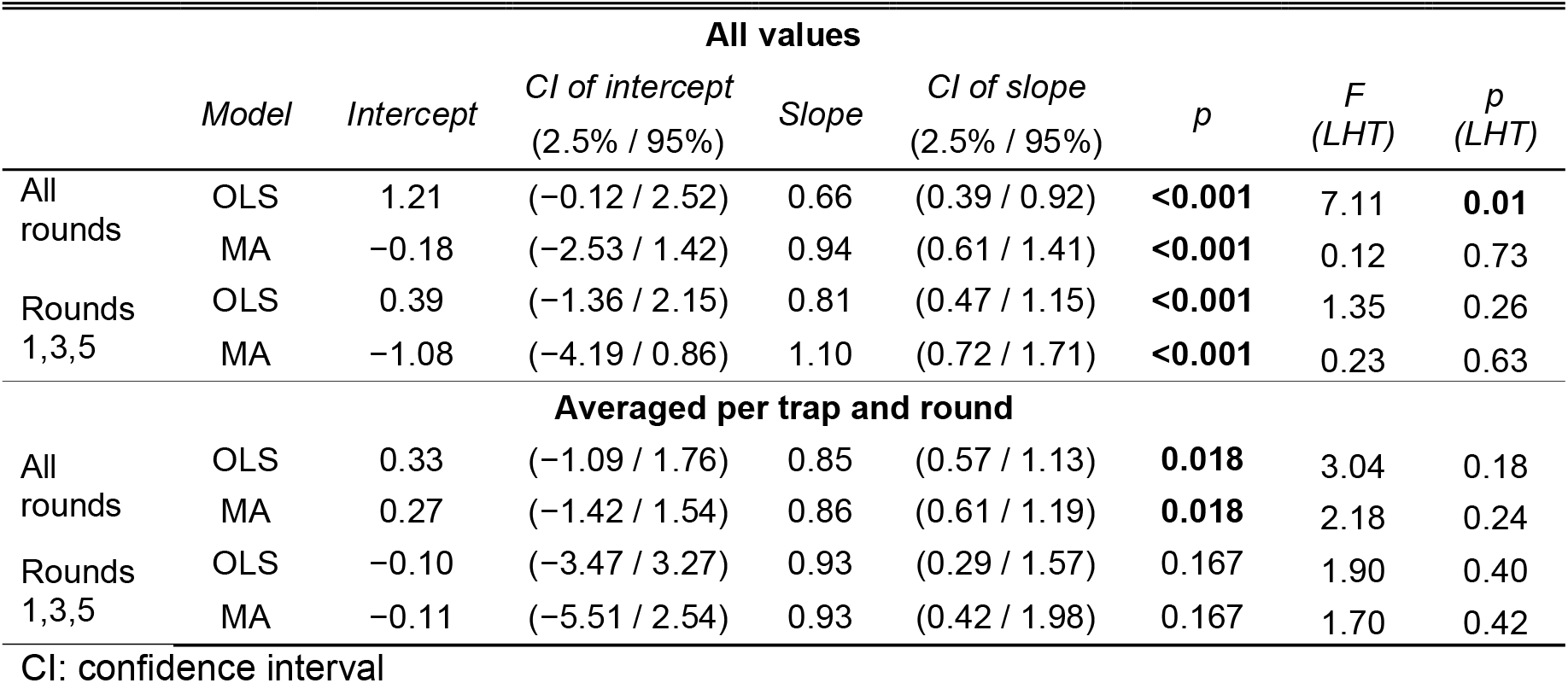
Results of the ordinary least squares (OLS) and major axis (MA) regressions of the number of individuals captured by the automated moth trap (AMT) and by a conventional light trap based on a complete (all rounds) and a reduced (rounds 1,3,5) dataset. The results of a linear hypothesis test (LHT), testing whether slopes deviate from one, are also displayed. Abundance values were log–transformed prior to the analyses.

Sampling conducted later in the year resulted in fewer recorded individuals overall, independent of whether Round, as an ordered factor (see Appendix S6), or the Julian day, as an independent variable (Table 2 and Appendix Table S6.2.), was used. With Round, the average number of individuals (mean ± SE: 131 ± 24) captured per AMT was lower than the average number captured using the conventional traps (218 ± 36 individuals) (Appendix Table S6.3 and S6.4). This lower average was not apparent when the Julian day was used (Table 2 and Appendix S6.2). Nonetheless, the two trap types did not differ in depicting the seasonal decline of moths in the study area (Figure 2).

**Figure 2:**
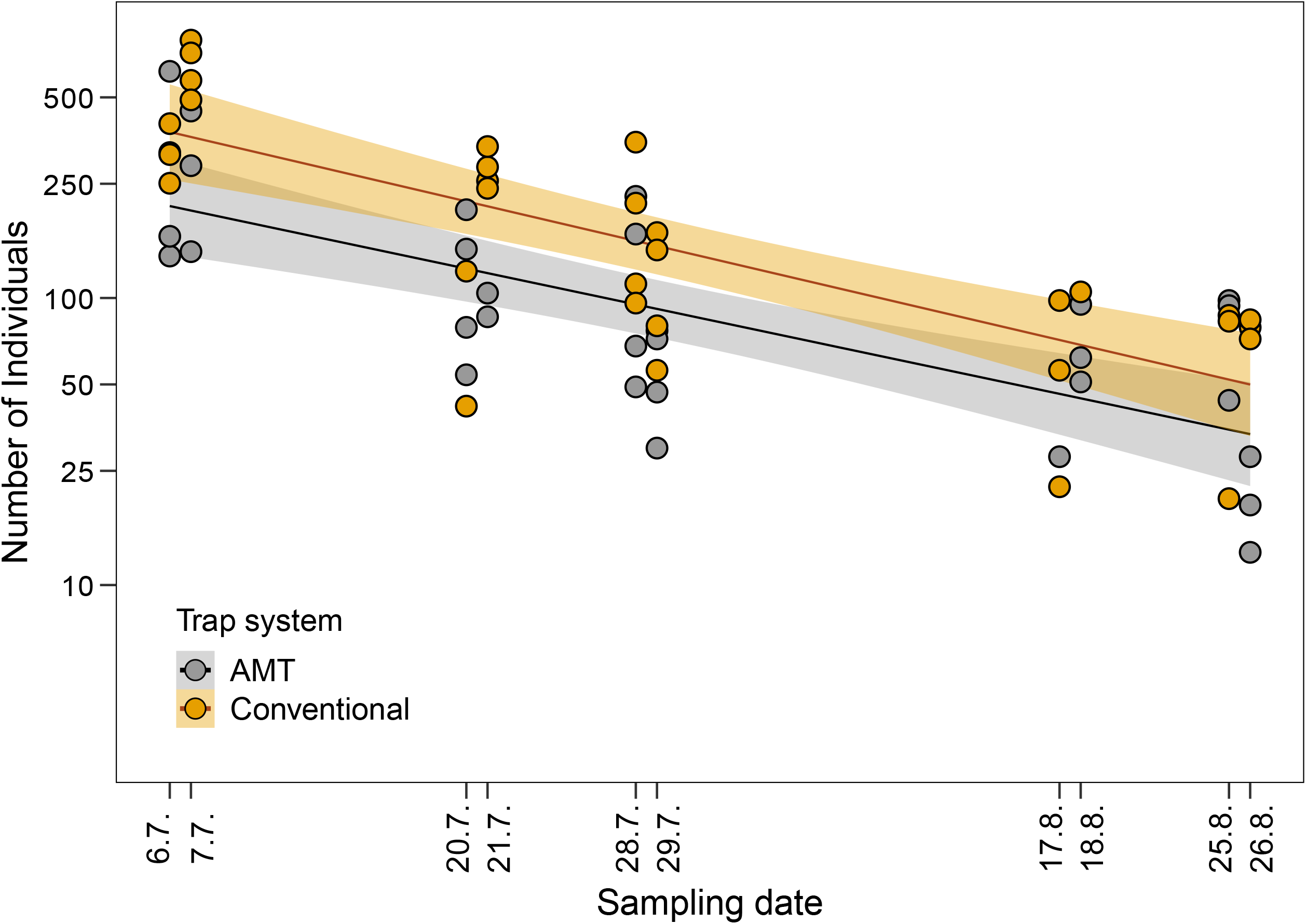
The number of individuals captured by all AMTs (grey) and by conventional light traps (orange) per sampling date. The positions of the data points indicate the Julian day of the sampling date. The shaded areas show the confidence intervals of the OLS regression. Only balanced data are shown. Note that the y-axis was log-transformed.

**Table 2:**
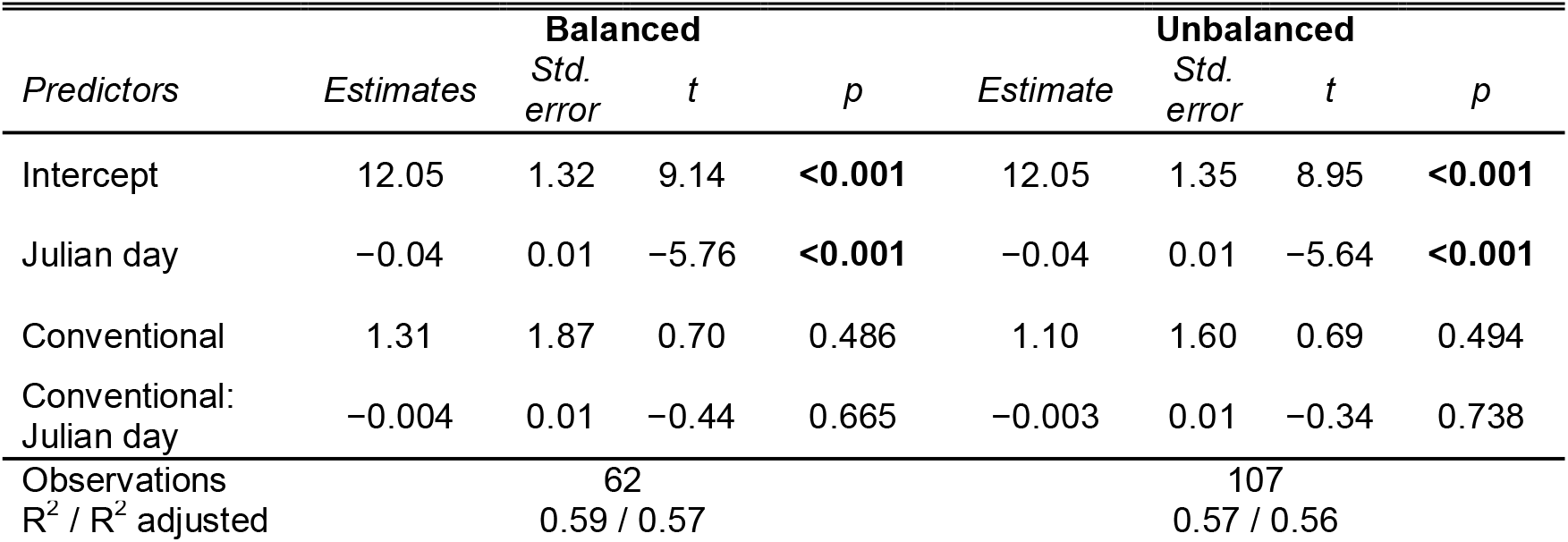
Results of the OLS regression testing potential differences in the trap systems with respect to seasonal trends in moth abundance, using the Julian day to account for differences in time intervals and the interaction with the trap type. The left column (Balanced) displays the results when only conventional traps and their corresponding AMTs were considered, and the right column (Unbalanced) when all conventional traps were included. Abundance values were log–transformed prior to the analyses.

| *Predictors* | *Estimates* | **Balanced** |  |  | *Estimate* | **Unbalanced** |  |  |
| --- | --- | --- | --- | --- | --- | --- | --- | --- |
|  |  | *Std. error* | *t* | *p* |  | *Std. error* | *t* | *p* |
| Intercept | 12.05 | 1.32 | 9.14 | **<0.001** | 12.05 | 1.35 | 8.95 | **<0.001** |
| Julian day | −0.04 | 0.01 | −5.76 | **<0.001** | −0.04 | 0.01 | −5.64 | **<0.001** |
| Conventional | 1.31 | 1.87 | 0.70 | 0.486 | 1.10 | 1.60 | 0.69 | 0.494 |
| Conventional:<br>Julian day | −0.004 | 0.01 | −0.44 | 0.665 | −0.003 | 0.01 | −0.34 | 0.738 |
| Observations |  | 62 |  |  |  | 107 |  |  |
| R<sup>2</sup> / R<sup>2</sup> adjusted |  | 0.59 / 0.57 |  |  |  | 0.57 / 0.56 |  |  |

## Discussion

Automation can facilitate the monitoring of insects at high temporal and spatial resolution, but the results must be representative (Collett & Fisher, 2017). Our AMT satisfies this criterion. While its light source differed from that of the conventional trap and did not operate continuously from dusk till dawn, the difference between the AMT and the conventional trap was negligible (and non-significant) with respect to seasonal effects. Overall, the AMT did not significantly differ from the conventional bucket trap in its ability to capture the temporal decline in the number of moths.

It should be noted that the number of individuals estimated using the AMT may have been too high, since some moths may have flown away during the 10 min without light and then returned to the screen after the light had been switched on again, such that they were counted several times. Conversely, counting moths without considering individual identities can lead to an underestimation of the total number, such as if one moth replaces another between frames. Most insect monitoring methods measure activity rates rather than abundance (Didham et al., 2020), as would have been the case with our AMT, because individuals may have visited the traps multiple times. By contrast, in conventional monitoring systems they would have been removed, thus potentially resulting in a depletion effect (Didham et al., 2020). Both trap types will be biased by environmental conditions (Appendix 7.2), such as the effect of temperature on activity (Wölfling et al., 2016). However, the AMT provides additional information on individuals arriving throughout the night, whereas only aggregated information is possible with conventional traps. The temporal data generated using the AMT allows us to address the effect of these environmental conditions with higher accuracy and detail (Appendix S7.1). Taken together, the AMT and bucket traps were similar in terms of the variance within their captured abundances despite the possible sources of uncertainty. Thus, our study showed that the AMT is sufficiently robust to capture important trends in moth abundance.

Off-the-shelf systems, especially those that are self-built and open-source, can help to broaden the reach of a monitoring campaign. Their advantages include easy access to technology, reproducibility, cost efficiency, increased data quality and quantity, the coverage of larger areas, and the incorporation of citizen science projects (Berger-Tal & Lahoz-Monfort, 2018). To achieve the goal of large-scale monitoring operations, knowledge sharing in the form of communication between working groups, such as via open source hardware and software designs, is vital to avoid unnecessarily redundant working steps during project development. Moreover, complex system-assembly procedures should be kept as simple as possible (and thoroughly tested in the field) to ensure end-user friendliness (Hill et al., 2019). Broader research community implementation and long term support for a monitoring system can also prolong its use, such as by offering solutions to malfunctions (Hill et al., 2019). Thus, automated monitoring systems, when coupled with technological innovations in data analytics and automated data validation, can form the basis of conservation-relevant decisions (August et al., 2015).

### Field-work applicability and lessons learned

Preti et al. (2021) summarized several requirements for efficient camera monitoring systems. Although these systems were originally devised for monitoring pests, many of their design aspects also apply to camera systems for monitoring biodiversity. The requirements included: user-friendly, easy to maintain, cost efficient, long operation times, low energy consumption, and sufficiently robust to withstand environmental conditions such as rain, dew, or wide temperature ranges. Camera-based systems that fulfill these criteria support scalability, i.e., monitoring at high spatial and temporal resolution. However, the first step in developing a new biodiversity monitoring system is to obtain a successful prototype, but its in-field applicability, handleability or the ability to depict ecological patterns with satisfactory data quality is often neglected. Among the goals in developing our adapted AMT was to create a system that was convenient, waterproof, and both cost and energy-efficient.

Extensive in-field use of our ATM highlighted two issues that need to be addressed. First, the DIY prototypes constructed from consumer-class hardware suffered problems in the field, despite extensive prior laboratory testing, which resulted in data loss (Appendix S5). These problems included the loosening of soldered connections during transport over rough terrain, corrupted SD cards, and modifications/undocumented changes by the manufacturer in the specifications of individual components that led to incompatibilities with the Raspberry Pi. These technical failures could only be identified during field work, but they can be fixed inexpensively.

The second problem was related to maintenance issues. Most of the maintenance of the boxes involved changing power banks and SD cards. The latter allowed faster data transfer than achieved with WLAN. In this study, the power supply of the AMTs consisted of portable energy storage, since the number of AMTs was limited and the traps had to be shared between plots. In addition, our criteria for the AMT included a setup that recycled many of the resources used in conventional light traps and that avoided the need for additional infrastructure, such as a power supply from solar panels. However, in a long-term deployment the additional investment in photovoltaic systems or, in shadier locations, shade-tolerant solar panels, would be worthwhile. A similarly constructed bat monitoring system, described in a previous study (Gottwald et al., 2021), could likewise be operated autonomously. For data transfer, rather than frequent visits to the trap sites to exchange the SD card, the research site could be remotely accessed. Data could be directly extracted using LTE sticks (Gottwald et al., 2021), or, when restricted by mobile phone reception, LoRa (Ferreira et al., 2020), which would again require added infrastructure. A further necessity is data-processing software (a trained neural network) that would allow the Raspberry Pi to analyze and process the images in-field, thus reducing the size of the output files.

Our tested AMT setup had total material costs of 511 € per trap, not including their working time. The cost of a conventional trap is 240 €, with additional running costs of 4 € per night for the chloroform. While the AMT is currently more expensive, over the long-term its cost will be compensated by the savings related to its operational advantages, especially regarding personnel costs and the need for frequent visits to the AMT.

### Potential of AMTs

Our primary interest was to determine whether the AMT can provide information on moth abundances; we were not interested in species identification yet. The system works with all cameras that use a ribbon cable and can be controlled by the Raspberry Pi; it also uses any 5 V light source that does not need pulse duration modulation. With a better camera and better illumination, the image resolution may be sufficiently high to at least allow the identification of conspicuous species or even the tracking of individuals (Bjerge et al., 2021; Korsch et al., 2021). However, AMTs might differ from conventional traps in their effect on species behavior, resulting in differences in the composition of the recorded species (Brehm & Axmacher, 2006; Wölfling et al., 2016). Further studies should test for differences in the species composition obtained with the AMT and conventional traps.

The manual identification of species using conventional traps can be time-consuming and cumbersome. This was highlighted by Hausmann et al (2020), since 9 years after insects were collected using malaise traps (Ssymank & Doczkal, 2017), only 10% of the collected individuals had been identified. Although methods such as DNA metabarcoding from pooled samples could accelerate species identification, they result in a considerable loss of information on abundance (Ji et al., 2013). The further combination with automated species identification tools, e.g., iNaturalist (2021), whose capabilities are constantly expanding and improving (Korsch et al., 2021), would be beneficial. Such tools make use of machine learning algorithms such as artificial neural networks, which after sufficient training can recognize and distinguish nearly all specimens with an accuracy comparable to that of humans (Ajit et al., 2020). A manual count of individuals from AMT images does not save time compared to conventional insect sorting, since depending on the activity rates of the insects, it may take up to an hour per trap and night to count the moth individuals depicted in the approximately 1500 AMT images accumulated in a single night. Moreover, insect movement in consecutive images could make it difficult to assess the momentary insect count or to register moth arrivals and departures. While the sorting of sampled insects from conventional traps is likely to be considerably shorter, automated object detection using trained neural networks would make automated identification nearly instantaneous.

We have not yet tested the effects of different light schedules, but shorter UV and LED screen illumination time would save energy; for example, a reduction to 30 min would save ~7 Wh (see Appendix S2). Moreover, it would limit disturbance of the moths (and other animals) and thus address an ethical issue in moth monitoring, i.e., the continuous operation of light traps, especially in sensitive regions, despite several studies reporting the negative effects of light pollution on moths (Boyes et al., 2021; Degen et al., 2016). Although a shorter light period could lead to a loss of information, this could be compensated by the use of several short light periods. Compared to a single light period lasting several hours, this strategy may better capture changes in species occurrences throughout the night (Scalercio et al., 2009). Indeed, as in Bjerge (2021), we observed changes in peak abundance throughout the sampling period (see Appendix S7). Different settings should be tested in further studies to find a reasonable compromise between a comprehensive assessment of night-flying insect fauna and the avoidance of excess disturbance, by keeping the light periods as short as possible.

Finally, in addition to moths, other light-attracted insects could be monitored in a similar, non-lethal, fashion using AMTs. One such candidate is adult caddisflies, an important component of macrozoobenthos communities in the vicinity of streams and rivers. Since their abundance mirrors the ecological conditions of lotic freshwater (Stuijfzand et al., 1999), caddisflies have been included in measures established by European environmental programs (Directive 2000/60/EC for water policy).

### Conclusion

Every design change to a trap, whether automated or not, can affect capture rates and lead to trap-dependent variability (Burner et al., 2021). Accordingly, data generated by a newly designed trap must be compared to those obtained with conventional traps. This study provided a first demonstration of the comparability of an inexpensive AMT with a conventional trap in capturing moth abundance trends. Our AMT can be built from low-cost DIY components and is capable of tracking changes in moth abundance. Nonetheless, the road from prototype to a ready-to-use monitoring system is paved with many challenges, including the decision whether to invest in energy-providing infrastructure that supports data transfer or to accept a lower resolution of the data to save on short-term costs. The choice will ultimately depend on the conditions at the respective sites and the funding framework of the monitoring program. With its modularity and flexible light scheduling, the AMT can be used to investigate several research questions while also reducing the disturbance of non-target animals in the vicinity of the trap. Perhaps most importantly, the AMT allows for high temporal resolution and long-term application in the field, without the ethical issues associated with conventional moth sampling.

## Acknowledgments

The research was funded by the Hessen State Ministry for Higher Education, Research and the Arts, Germany, as part of the LOEWE priority project Nature 4.0 – Sensing Biodiversity (https://uni-marburg.de/natur40). Moth sampling was permitted by the Lower Nature Conversation Authority Marburg and the Department of Nature Conversation of the district Marburg-Biedenkopf. We thank Hendrik Reers from OekoFor for sharing his experiences with the implementation of AMTs. Special thanks to Wendy Ran for copy-editing the manuscript.

## Conflict of interest

The authors declare no conflict of interest.

## Data availability

The operating system for the AMT can be downloaded from our Github repository (https://github.com/Nature40/InsectPhotoTrapOS/releases/tag/IPTv1.0.7). Datasets will be archived via data_UMR, the long-term data repository of the University of Marburg.

## Author contributions

L.H., N.F., J.M.M., and P.L. designed the study and build the traps. P.L. developed the software based on previous work with J.G. Field work was conducted by J.M.M. and L.H. with the help from M.F. The traps were maintained under supervision of P.L. Statistical analyses were performed by L.H. and N.F., image analyses were performed by J.M.M. Supporting analyses were performed by M.F. Guidance throughout the project was given by N.F., R.B., M.B., B.F. and T.N. The authors J.M.M., P.L. and L.H. contributed equally to the writing of the manuscript. All co-authors gave significant input and feedback on the manuscript.

## Notes

### Competing Interest Statement

The authors have declared no competing interest.

