## Supplement for "Automated non-lethal moth traps can be used for robust estimates of moth abundance"

*Shared first author

Corresponding author:

Jonas Mielke Möglich

#### S1) Moth traps

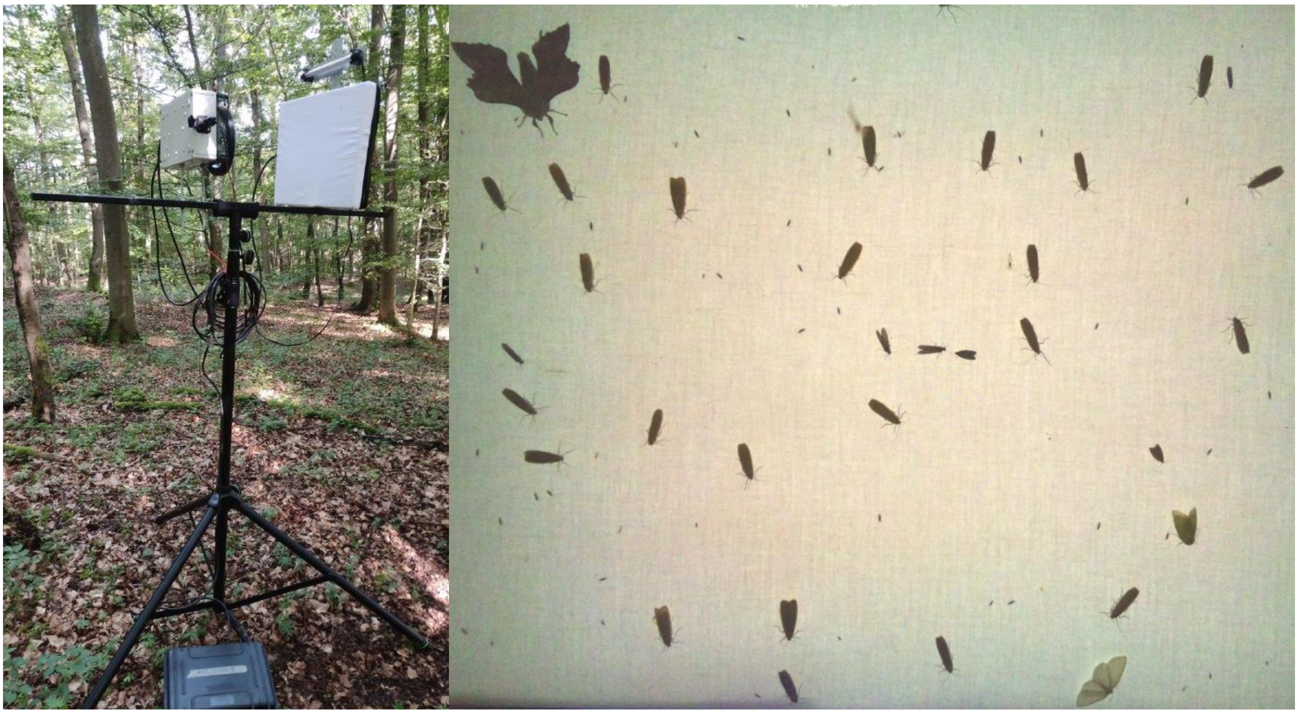

**Figure S1.1:** Photo of our automated moth trap (AMT) in the field setup and example of a photo taken by our AMT.

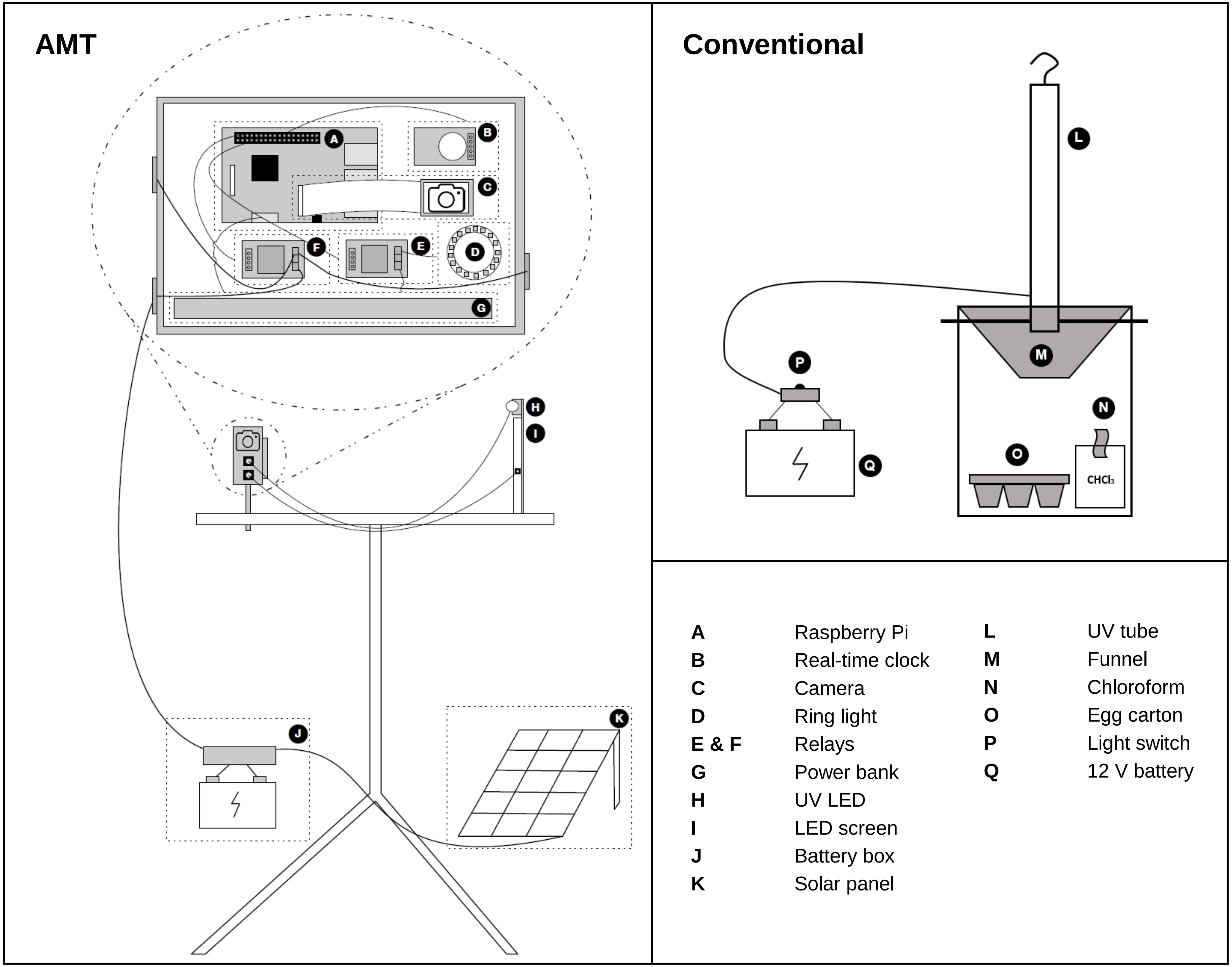


**Figure S1.2:** Architecture of our AMT setup and of a conventional bucket light trap.

#### S2) Power consumption

Power consumption was measured to provide an overview of how long the ATM can be operated using different energy sources:

- UV LED: 0.8A@12.5V ~ 10W
- LED Board: 0.78A@12.5V ~ 9.75W
- Ring light: limited by the output of the power bank
- Power bank: 2.1A@5.1V ~ 11W
- Raspberry Pi: 0.35A@5.1V ~ 1.8W

The used energy is the sum of the energy used in moth attraction multiplied by the percentage of attractions per hour and the energy consumption per photo multiplied by the number of photos taken in one hour and the power consumption of the Raspberry Pi.

The energy per hour is calculated as follows:

Energy use per h = t * (uv + led) + (n * s * rl / 3600) + rp

50 min attraction: 50/60 * (10 + 9.75) + (100 * 3 * 11 / 3600) + 1.8 = 19.175 Wh

40 min attraction: 40/60 * (10 + 9.75) + (80 * 3 * 11 / 3600) + 1.8 = 15.7 Wh

30 min attraction: 30/60 * (10 + 9.75) + (60 * 3 * 11 / 3600) + 1.8 = 12.225 Wh

20 min attraction: 20/60 * (10 + 9.75) + (40 * 3 * 11 / 3600) + 1.8 = 8.75 Wh

10 min attraction: 10/60 * (10 + 9.75) + (20 * 3 * 11 / 3600) + 1.8 = 5.275 Wh

where

t is the percentage of attraction, n is photos per hour, s is the duration of ring light per photo, uv is Watts of UV light, led is Watts of LED board , rl is watts of ring light, and rp is Watts used by the Raspberry Pi

### S3) Price list of AMT components

| | Modul | Parts (exemplary products) | Quantity | Price [€] (29.11.21) | Price per trap [€] |
|---|---|---|---|---|---|
| A1 | Camera box | Raspberry Pi 3 B | 1 | 59.99 | 59.99 |
| A2 | Camera box | SD card 64Gb | 1 | 15.99 | 15.99 |
| A3 | Camera box | Case | 1 | 23.00 | 23.00 |
| A4 | Camera box | Mounting plate (3d print) | 1 | 1.00 | 1.00 |
| A5 | Camera box | Jumper wires | 0.1 | 5.35 | 0.54 |
| A6 | Camera box | Neutrik NAC3FPX power socket female | 2 | 8.26 | 16.52 |
| A6 | Camera box | Neutrik NAC3MPX power socket male | 1 | 3.66 | 3.66 |
| A6 | Camera box | Neutrik SCNAC-FPX | 3 | 1.08 | 3.24 |
| B | Camera box | DS3231 RTC Modul I2C real-time clock AT24C32 for Arduino; with LIR2032 battery | 1 | 5.00 | 5.00 |
| C | Camera box | Raspberry Pi camera | 1 | 26.04 | 26.04 |
| D | Camera box | LED ring light | 1 | 8.99 | 8.99 |
| E,F | Camera box | Relay KY019RM | 2 | 3.79 | 7.58 |
| | | Subtotal camera box | | | 171.55 |
| G | Power | Power bank | 1 | 47.00 | 47.00 |
| H | Light box | UV light | 1 | 33.00 | 33.00 |
| I1 | Light box | Box for backlight | 1 | 23.30 | 23.30 |
| I2 | Light box | LED strip for backlight | 0.4 | 16.44 | 6.58 |
| I3 | Light box | Aluminum foil in format of box for backlight | 0.2 | 3.00 | 0.60 |
| I4 | Light box | Lee filter in format of box for backlight | 0.03 | 65.00 | 1.95 |
| I5 | Light box | Bedsheet in format of box for backlight | 0.1 | 10.00 | 1.00 |
| I6 | Light box | Spray adhesive | 0.2 | 15.00 | 3.00 |
| I7 | Light box | Neutrik NAC3MPX power socket male | 1 | 3.66 | 3.66 |
| I7 | Light box | Neutrik SCNAC-FPX | 1 | 1.08 | 1.08 |
| I8 | Light box | Neutrik NAC3MX-W plug male | 1 | 7.44 | 7.44 |
| I9 | Light box | Cable connections | 1 | | 1.00 |
| | | Subtotal light box | | | 129.61 |
| J1 | Power | Battery | 1 | 32.18 | 32.18 |
| J2 | Power | Battery box | 0.5 | 52.80 | 26.40 |
| J3 | Power | Neutrik NAC3FPX power socket female | 1 | 8.26 | 8.26 |
| J3 | Power | Neutrik SCNAC-FPX | 1 | 1.08 | 1.08 |
| | Power | Fuse | 1 | 0.50 | 0.50 |
| | | Subtotal power box | | | 68.42 |
| K | Cable | Neutrik NAC3FX-W plug female | 2 | 9.08 | 18.16 |
| K | Cable | Neutrik NAC3MX-W plug male | 2 | 7.44 | 14.88 |
| K | Cable | Cable | 1 | | 2.00 |
| | | Subtotal cable | | | 35.04 |

| | | | | |
|---|---|---|---|---|
| Mounting | Backlight and UV light holder self-made | 1 | 9.30 | 9.30 |
| Mounting | Tripod | 1 | 72.00 | 72.00 |
| Mounting | Threaded rods | 1 | 1.00 | 1.00 |
| Mounting | Screws | 4 | 1.00 | 4.00 |
| | Subtotal mounting | | | 86.30 |
| | Other items | | | 20.00 |
| **Total** | | | | **510.92** |

#### S4) Software

Exemplary configuration of our test unit and sample runs:

*[TimedInsectTrapAnalysesUnit]*

*light_pin = 26*

*uv_light_pin = 21*

*hourly = True*

*uv_light_start_time = 00*

*uv_light_end_time = 50*

*adjust_time_in_seconds = 2*

*take_photo_every_seconds = 30*

*[run.1]*

*start = 21:10*

*stop = 23:59:59*

*[run.2]*

*start = 00:00*

*stop = 05:50*

[TimedInsectTrapAnalysesUnit] defines the unit itself and the associated parameters, including the pins used for the respective relays as well as the start and end times of the hourly routine. These values can be adapted as needed, included depending on the unit and the research question. The time the camera needs to focus is specified by adjust_time_in_seconds: for a camera with a fixed focus, this option can be set to 0. The time between two photos is specified by take_photo_every_seconds, for photos taken periodically between uv_light_start_time and uv_light_end_time.

To map as many use cases as possible, different runs can be defined and the parameters modified in each one. For example, take_photo_every_seconds can be set to 30 in run 1 and to 60 in run 2.

#### S5) Test of the effect of location and sampling date

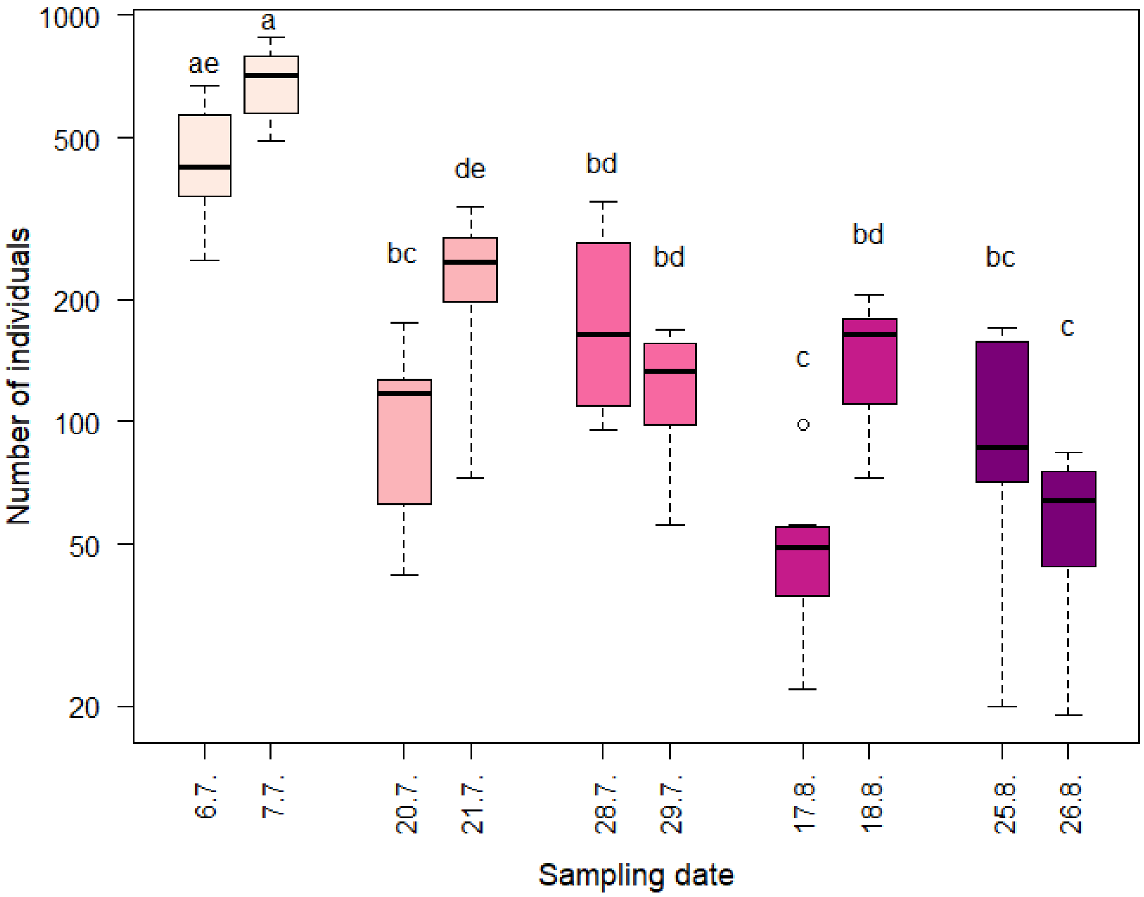


**Figure S5.1:** Results of the ANOVA and subsequent Tukey HSD to test for pairwise differences in the sampling dates. Boxes are color-coded according to the sampling round. Pairs that are not significantly different from each other share the same label. Round 2, sampled on 20$^{th}$ and 21$^{st}$ July, and round 4, sampled on 17$^{th}$ and 18$^{th}$ August, differ from each other; round 1, sampled on 6$^{th}$ and 7$^{th}$ July, round 3, sampled on 28$^{th}$ and 29$^{th}$ July, and round 5, sampled on 25$^{th}$ and 26$^{th}$ August, do not (p-values 0.001 and 0.025, respectively).

ANOVAs and subsequent post hoc tests were used to determine whether the localities or dates differed from each other with respect to the number of moths caught by the conventional traps. Prior to the analysis, the number of individuals was log-transformed. The number of moths differed between dates ($F_{9,66}$ = 26.47, $p < 0.001$; Figure S5.1) but not between the plots (ANOVA, $F_{15,60}$ = 0.42, p = 0.967). Tukeys HSD test showed that the two subsequently sampled dates differed in round 2 (p = 0.02, 95 % CI = [0.06, 1.66]) and round 4 (p = 0.001, 95 % CI =

[0.29, 1.88]), but not in round 1 (p = 0.65, 95 % CI = [-0.32, 1.23]), round 3 (p = 0.85, 95 % CI = [-1.14, 0.40]), or round 5 (p = 0.62, 95 % CI = [-1.32, 0.32]).

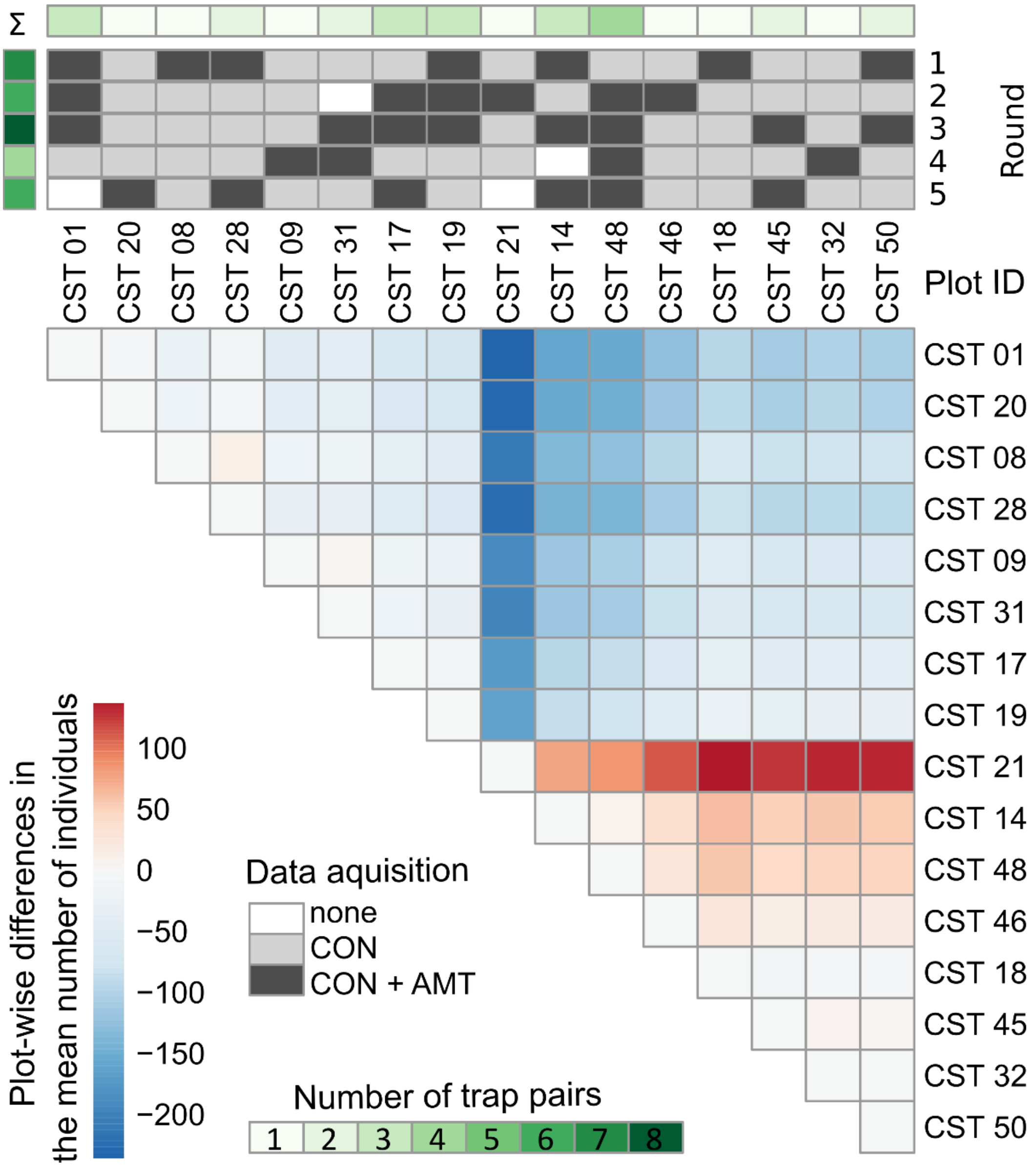


**Figure S5.2:** Overview of data acquisition per plot and sampling rounds using conventional light traps (CON) and our automated moth traps (AMT) (top) and plot-wise differences in the mean number of individuals, determined using the package "pheatmap" (Kolde, 2019). Only individuals sampled from CON traps were used for calculation of plot-wise differences. Differences are indicated along a color scale from red (positive) to blue (negative). None of the differences were significant. The top rows indicate whether the AMT acquired data during the respective round (dark grey) or whether data was lost.


Kolde, R. (2019). *pheatmap: Pretty Heatmaps* (1.0.12) [Computer software]. https://CRAN.R-project.org/package=pheatmap

#### S6) Main analysis in the form of generalized linear models (GLMs) with negative binomial distribution

**Table S6.1:** Results of GLM modeling the number of individuals captured by the automated moth traps (AMTs) based on the (log-transformed) number of moths caught by the conventional traps. Numbers in parentheses refer to the confidence intervals. The left column reports the results when all rounds were considered, and the right column when only those rounds (1, 3, 5) with comparable consecutive nights were included.

| | **All rounds** | | | | **Rounds 1,3,5** | | | |
|---|---|---|---|---|---|---|---|---|
| *Predictor* | *Log-mean* | *Std. error* | *z* | *p* | *Log-mean* | *Std. error* | *z* | *p* |
| Intercept | 1.31 (0.25 – 2.42) | 0.59 | 2.25 | **0.025** | 0.46 (-0.98 – 1.97) | 0.74 | 0.63 | 0.530 |
| log (conventional) | 0.67 (0.46 – 0.89) | 0.12 | 5.85 | **<0.001** | 0.84 (0.55 – 1.12) | 0.14 | 5.89 | **<0.001** |
| Observations | 31 | | | | 21 | | | |
| $R^2$ Nagelkerke | 0.78 | | | | 0.85 | | | |

**Table S6.2:** Results of GLM modeling the number of individuals vs. the Julian day in the interaction with trap type. Numbers in parentheses refer to the confidence intervals. The left column (Balanced) reports the results when only the conventional traps with corresponding AMTs were considered, and the right column (Unbalanced) the results when all conventional traps were included.

| | **Balanced** | | | | **Unbalanced** | | | |
|---|---|---|---|---|---|---|---|---|
| *Predictor* | *Mean* | *Std. error* | *z* | *p* | *Mean* | *Std. error* | *z* | *p* |
| Intercept | 12.01 (9.84 – 14.21) | 1.19 | 10.09 | **<0.001** | 12.01 (9.75 – 14.30) | 1.24 | 9.68 | **<0.001** |
| Julian day | −0.03 (−0.05 – −0.02) | 0.01 | −6.18 | **<0.001** | −0.03 (−0.05 – −0.02) | 0.01 | −5.93 | **<0.001** |
| Conventional | 1.47 (−1.64 – 4.58) | 1.69 | 0.87 | 0.382 | 1.06 (−1.63 – 3.74) | 1.47 | 0.72 | 0.47 |
| Conventional: Julian day | −0.002 (−0.02 – 0.01) | 0.01 | −0.60 | 0.549 | −0.002 (−0.02 – 0.01) | 0.01 | −0.35 | 0.729 |
| Observations | 62 | | | | 107 | | | |
| $R^2$ Nagelkerke | 0.87 | | | | 0.85 | | | |

**Table S6.3:** GLM modeling the number of individuals vs. the round (Round) as an ordered factor in the interaction with trap type. Numbers in parentheses refer to the confidence intervals. The left column (Balanced) reports the results when only conventional traps with a corresponding AMT were considered, and the right column (Unbalanced) when the results of all conventional traps were included.

| | **Balanced** | | | | **Unbalanced** | | | |
|---|---|---|---|---|---|---|---|---|
| *Predictors* | *Log-mean* | *Std. error* | *z* | *p* | *Log-mean* | *Std. error* | *z* | *p* |
| Intercept | 4.59<br>(4.40 – 4.79) | 0.10 | 46.34 | **<0.001** | 4.59<br>(4.40 – 4.79) | 0.10 | 46.11 | **<0.001** |
| Round [linear] | −1.35<br>(−1.78 – −0.92) | 0.22 | −6.20 | **<0.001** | −1.35<br>(−1.78 – −0.92) | 0.22 | −6.17 | **<0.001** |
| Round [quadratic] | 0.37<br>(−0.04 – 0.79) | 0.21 | 1.76 | 0.079 | 0.37<br>(−0.04 – 0.79) | 0.21 | 1.75 | 0.080 |
| Round [cubic] | −0.17<br>(−0.65 – 0.30) | 0.24 | −0.70 | 0.484 | −0.17<br>(−0.65 – 0.30) | 0.24 | −0.70 | 0.486 |
| Round [4$^{th}$ degree] | 0.19<br>(−0.25 – 0.61) | 0.22 | 0.85 | 0.394 | 0.19<br>(−0.25 – 0.61) | 0.22 | 0.85 | 0.396 |
| Conventional | 0.44<br>(0.17 – 0.71) | 0.14 | 3.16 | **0.002** | 0.52<br>(0.29 – 0.75) | 0.12 | 4.49 | **<0.001** |
| Conventional:<br>Round [linear] | −0.24<br>(−0.85 – 0.36) | 0.31 | −0.79 | 0.428 | −0.09<br>(−0.60 – 0.42) | 0.26 | −0.35 | 0.730 |
| Conventional:<br>Round [quadratic] | −0.02<br>(−0.61 – 0.56) | 0.30 | −0.08 | 0.935 | 0.09<br>(−0.40 – 0.58) | 0.25 | 0.36 | 0.716 |
| Conventional:<br>Round [cubic] | 0.25<br>(−0.41 – 0.91) | 0.34 | 0.75 | 0.456 | −0.08<br>(−0.62 – 0.47) | 0.28 | −0.30 | 0.761 |
| Conventional:<br>Round [4$^{th}$ degree] | 0.08<br>(−0.53 – 0.68) | 0.31 | 0.25 | 0.803 | 0.10<br>(−0.40 – 0.61) | 0.26 | 0.39 | 0.700 |
| Observations | 62 | | | | 107 | | | |
| $R^2$ Nagelkerke | 0.91 | | | | 0.92 | | | |

**Table S6.4**. OLS (linear model) results of the number of individuals vs. Round as an ordered factor and independent variable. Numbers in parentheses refer to the confidence intervals. The left column (Balanced) reports the results when only the conventional traps with a corresponding AMT were considered, and the right column (Unbalanced) when all conventional traps were considered. Note that the values were log-transformed.

| | **Balanced** | | | | **Unbalanced** | | | |
|---|---|---|---|---|---|---|---|---|
| *Predictor* | *Estimates* | *Std. error* | *t* | *p* | *Estimates* | *Std. error* | *t* | *p* |
| Intercept | 4.42 (4.19 – 4.66) | 0.11 | 38.48 | **<0.001** | 4.42 (4.21 – 4.64) | 0.11 | 40.20 | **<0.001** |
| Round [linear] | −1.43 (−1.94 – −0.92) | 0.25 | −5.65 | **<0.001** | −1.43 (−1.91 – −0.95) | 0.24 | −5.90 | **<0.001** |
| Round [quadratic] | 0.31 (−0.18 – 0.80) | 0.24 | 1.25 | 0.216 | 0.31 (−0.16 – 0.77) | 0.23 | 1.31 | 0.194 |
| Round [cubic] | −0.21 (−0.76 – 0.35) | 0.28 | −0.75 | 0.455 | −0.21 (−0.74 – 0.32) | 0.27 | −0.79 | 0.433 |
| Round [$4^{th}$ degree] | 0.08 (−0.43 – 0.58) | 0.25 | 0.30 | 0.767 | 0.08 (−0.40 – 0.55) | 0.24 | 0.31 | 0.756 |
| Conventional | 0.47 (0.14 – 0.79) | 0.16 | 2.87 | **0.006** | 0.54 (0.28 – 0.80) | 0.13 | 4.16 | **<0.001** |
| Conventional: Round [linear] | −0.17 (−0.89 – 0.55) | 0.36 | −0.48 | 0.631 | −0.10 (−0.67 – 0.47) | 0.29 | −0.36 | 0.720 |
| Conventional: Round [quadratic] | 0.12 (−0.57 – 0.82) | 0.35 | 0.35 | 0.725 | 0.19 (−0.37 – 0.74) | 0.28 | 0.67 | 0.506 |
| Conventional: Round [cubic] | 0.27 (−0.52 – 1.05) | 0.39 | 0.68 | 0.499 | −0.07 (−0.68 – 0.54) | 0.31 | −0.23 | 0.822 |
| Conventional: Round [$4^{th}$ degree] | 0.23 (−0.49 – 0.94) | 0.36 | 0.63 | 0.529 | 0.29 (−0.28 – 0.85) | 0.29 | 1.00 | 0.318 |
| Observations | 62 | | | | 107 | | | |
| $R^2$ / $R^2$ adjusted | 0.63 / 0.57 | | | | 0.65 / 0.61 | | | |

#### S7) Moth individuals overnight

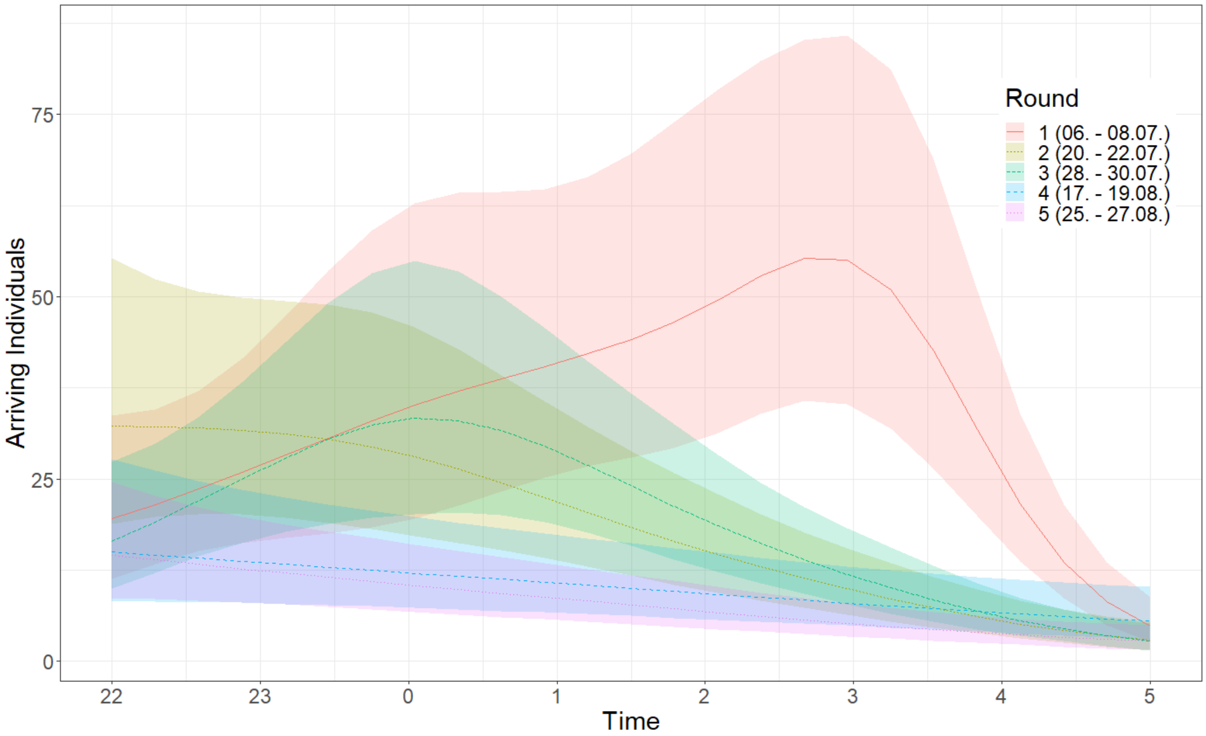


**Figure S7.1**: Generalized additive models (GAMs) (Coretta et al., 2021; Wood et al., 2016) of individuals arriving per time, listed by rounds. The round was inserted as a linear term; time was added using a cubic spline basis function, with $k = 5$. To account for the effect of the plot in the repeated measurements of arriving individuals, the trap location was included as a random effect. The colored lines represent the GAMs of the different rounds, and the shaded areas the confidence intervals. According to the model ${R^2}_{(adj.)} = 0.722$ , the largest number of individuals overall was captured during round one, i.e., the first and earliest of the rounds. Also, moth arrivals as recorded by the AMT in round one peaked at 3 a.m., whereas in the other rounds the peak occurred at 11 p.m. or 12 a.m.. Since data from sunset (around 9 p.m.) was missing, such that the earliest hour of moth activity was not included in the model, an even earlier peak may have been missed.

Coretta, S., van Rij, J., & Wieling, M. (2021). *tidymv: Tidy Model Visualisation for Generalised Additive Models* (3.2.1) [Computer software]. https://CRAN.R-project.org/package=tidymv

Wood, S. N., Pya, N., & Säfken, B. (2016). Smoothing Parameter and Model Selection for General Smooth Models. *Journal of the American Statistical Association*, *111*(516), 1548–1563. https://doi.org/10.1080/01621459.2016.1180986

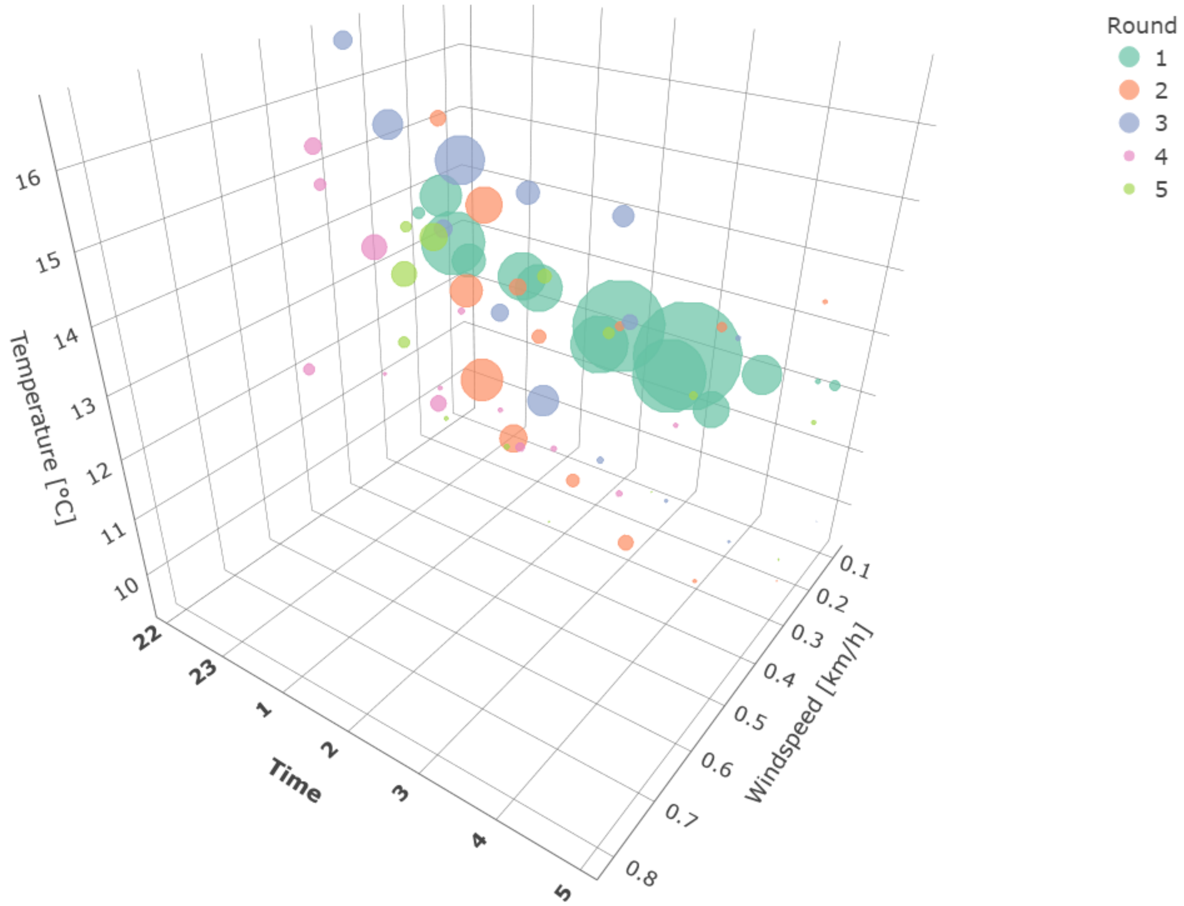


**Figure S7.2**: 3D scatter plot (Sievert, 2020) of recorded temperatures and wind speeds and the hours of data recording, beginning at 10 pm. The colors indicate the rounds; the sizes of the points shown are relative to the number of individuals counted. Thus, as seen in the plot, very large numbers of individuals were observed in the first round (large green dots), at lower windspeeds, medium temperatures, and late in the night (between 2 and 4 a.m.). The barely visible spots of later rounds indicate a very low number of moths arriving at the AMTs late at night, when temperatures were low and wind speeds were around 0.3 and 0.4 km/h.

Sievert, C. (2020). *Interactive Web-Based Data Visualization with R, plotly, and shiny*. CRC Press
